# Absolute scattering length density profile of liposome bilayers obtained by SAXS combined with GIXOS - a tool to determine model biomembrane structure

**DOI:** 10.1101/2022.12.13.520277

**Authors:** Chen Shen, Christian Woelk, Alexey G. Kikhney, Jaume Torres, Wahyu Surya, Richard D. Harvey, Gianluca Bello

## Abstract

Lipid membranes play an essential role in biology, acting as host matrices for biomolecules like proteins and facilitating their functions. Their structures, and structural responses to physiologically relevant interactions, i.e. with membrane proteins, provide key information for understanding biophysical mechanisms. Hence, there is a crucial need of methods to understand the effects of membrane host molecules on the lipid bilayer structure. Here, we present a purely experimental method for obtaining the absolute scattering length density (SLD) profile and the area per lipid of liposomal bilayers, by aiding the analysis of small angle X-ray scattering (SAXS) data with the volume of bare headgroups obtained from fast (20-120s) grazing incidence off-specular scattering (GIXOS) data from monolayers of the same model membrane lipid composition. The GIXOS data experimentally demonstrate that the variation of the bare headgroup volume upon lipid packing density change is small enough to allow its usage as a reference value without knowing the lipid packing stage in a bilayer. This approach also bares the advantage that the reference volume is obtained at the same aqueous environment as used for the model membrane bilayers. We demonstrate the validity of this method using several typical membrane compositions, as well as one example of a phospholipid membrane with an incorporated transmembrane peptide. This methodology allows to obtain absolute scale values rather than relative scale by using solely X-ray-based instrumentation, retaining a similar resolution of SAXS experiments. The presented method has high potential to understand structural effects of membrane proteins on the biomembrane structure.

## 1 Introduction

Lipid membranes are one of the structural foundations of living organisms. They support important biological activities, by acting as hosting matrices for embedded transporters and signal transducing receptors, as well as facilitating endocytosis and secretory processes. Their structures and modifications exert fundamental influences on these processes, and thus structural information is essential for understanding underlying mechanisms^1^. Whilst numerous methodological developments have improved the resolution of protein and peptide structures, comparable advances on their lipid matrices are less frequently made^2-5^, despite this information is of high interest to understand biological processes which depend on membrane bending, stiffening and thickness modulation. Determining high-resolution membrane structure under physiologically relevant conditions of milieu and lipid thermodynamics is equally important for understanding membrane biology at the molecular level.

Typical approaches in membrane studies use bilayer systems such as liposomes^2,5-7^, hydrated supported multilayers^7-10^, solid supported bilayers at bulk interfaces^11,12^, or monolayers at air-water or oil-water interfaces^4^. These models depict representative features of interest of more complex biological membranes, allowing high resolution structures to be acquired by applying suitable X-ray or neutron scattering methods^2,4-7,11,13^. Among these methods, small angle (X-ray) scattering (SAXS) studies on liposome dispersions of fully hydrated membranes are required to mimic biological settings. Such systems carry the advantage that the composition of the aqueous phase can be readily adjusted to model physiologically relevant conditions of pH, osmolarity, ionic strength or bulk ligand concentration.

SAXS data alone provides scattering length density (SLD) contrast profiles of phospholipid bilayers at sub-angstrom resolution^7^ which can clearly give some structural information of the model membrane structure along the membrane normal, such as the distance between the high SLD phosphate groups of the two leaflets, the thickness of the lower SLD hydrocarbon core, and the shape of the headgroup toward the bulk and towards the core region (example, see Figure 1A)^3,5^. However, this is not sufficient to provide other physiologically important bilayer parameters such as area and volume per lipid or the hydration state of the headgroups. Obtaining these parameters requires an absolute SLD profile in reciprocal area units, e.g. Å^-2^, to calculate the material density. Typically, SAXS data is obtained in arbitrary intensity units that reflect only the relative contrast variation from bulk solvent SLD. Measuring absolute SAXS intensity in reciprocal length units, e.g. cm^-1^, is rarely accurate enough, due to the difficulties in measuring transmission intensity and in knowing the exact number of scatterers in the volume illuminated by X-rays ^3,14^. An alternative approach is to use the bare headgroup volume, excluding hydration, obtained from membranes in the gel phase, as a reference value for systems with the same headgroup, as this volume remains independent from the membrane phase ^3,15^. In addition, they obtained the lipid specific volume of the SAXS sample in parallel with neutral buoyancy experiments^3,7,16^. Combining the two values yields the area per lipid and the scaling factor for the absolute SLD value. Later, simulation was also used to provide the specific volume of lipid in broader applications ^15^.

**Figure 1.**
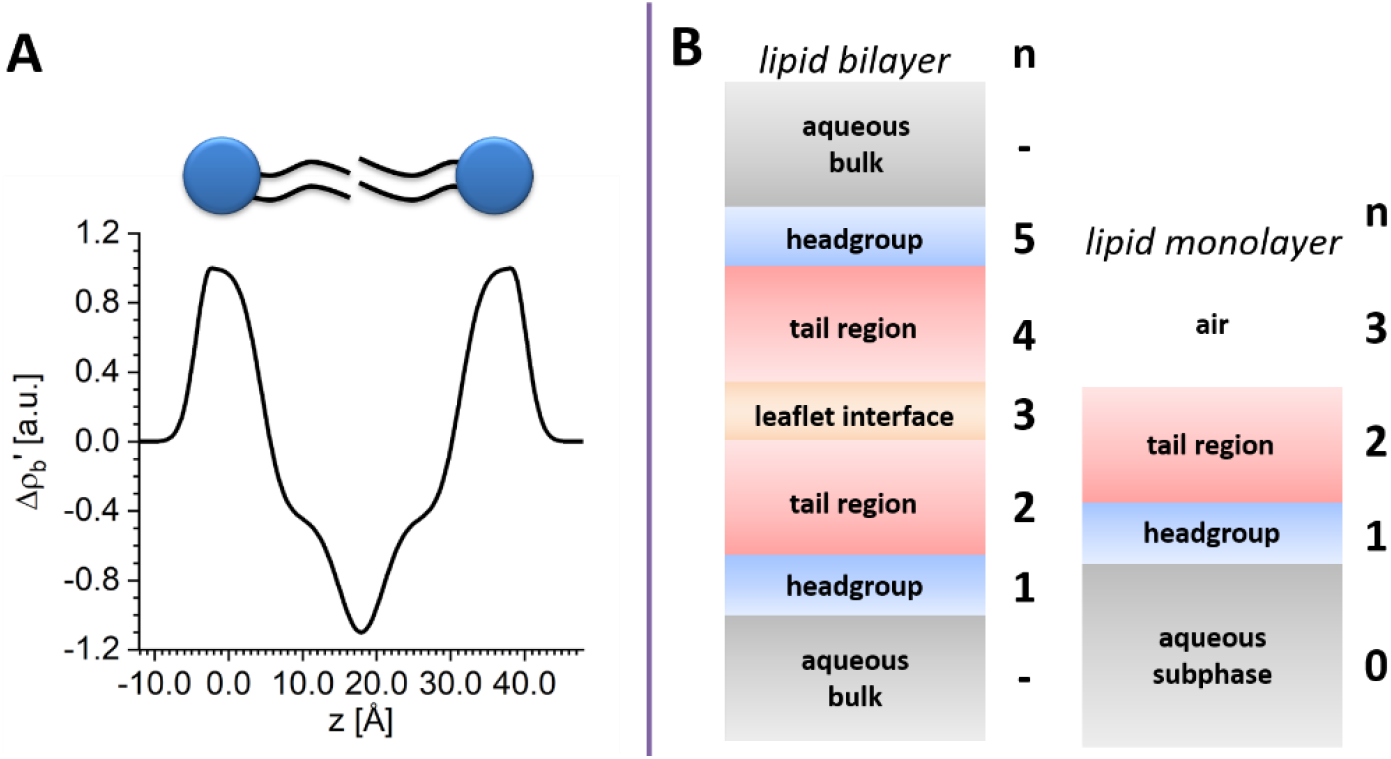
A) Example of a relative SLD contrast profile of a lipid bilayer in liposomes obtained direct from a SAXS data analysis. Distances (z) are determinable, but the SLD contrast 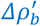 is given in arbitrary units. Obtaining absolute SLDs is not possible. B) Compartment models for a bilayer and for a monolayer.

Following the same logic, we present a new alternative volumetric approach for obtaining absolute SLD profiles of liposomes, requiring solely input from complementary, rapidly-acquired experimental X-ray data. Via the theoretical derivation, we show that the relative SLD contrast ratio within a lipid bilayer (hydrated head, tail) is sufficient to calculate the area per lipid, and, accordingly, the absolute SLD profile once the volume of the bare headgroup is known. Moreover, we present the bare headgroup volume determined experimentally from monolayers of the same composition on the same aqueous bulk at various lateral pressures using grazing incidence off-specular X-ray scattering (GIXOS)^17^. This GIXOS results demonstrate that variation of monolayer membrane phase and surface pressure has a negligible effect on the determination of absolute SLD profile and area per molecule in the bilayer from SAXS analysis. It justifies the use of this value as the input reference to analyze bilayer data, regardless of the phase state of the lipids.

The method is validated by examples of commonly used phosphatidyl choline (PC) liposomes and PC/phosphatidyl serine (PS) binary mixed liposomes, all unextruded, to enable better comparability with previous results. In addition, we include a PC/PS membrane with embedded α-helical, envelop (E) protein from the severe acute respiratory syndrome coronavirus 2 (SARS-CoV-2), a protein that is assumed to induce membrane curvature during virion formation^18^. This peptide-embedded membrane requires the modelling with an asymmetric bilayer profile, thus it validates a broader application of this approach than simple model systems. The presented results demonstrate the applicability of the method to understand model biomembranes, also in the presence of interacting molecules such as proteins and peptides.

## 2 Materials and Methods

### 2.1 Theoretical Section

Our approach to obtaining absolute SLD profiles consists of three steps. (I) An arbitrary scale SLD contrast profile is obtained from vesicle dispersions from a conventional analysis of SAXS data^19^. (II) Langmuir monolayers of the same lipid composition and aqueous solvent are measured at different surface pressures by GIXOS, up to 0.9 Å^-1^ in Q_z_, to obtain the monolayer absolute SLD profiles^17^. These monolayer SLD profiles provide the volume of the bare headgroup. (III) The absolute SLD profile of the bilayer is calculated from the SLD contrast profile by using the bare headgroup volume as the reference. In this section, the theoretical derivation for obtaining the absolute SLD profiles is presented first, followed by the experimental examples in the next two sections.

#### 2.1.1 Relative SLD contrast obtained from SAXS

The SAXS analysis we followed to get the arbitrarily scaled SLD profiles^3,20^, though being conventional, are briefly presented here for clarity. High resolution analysis of lipid bilayer structure from liposome suspensions requires the inclusion of at least three oscillations of a bilayer form factor, to roughly 0.6 Å^-1 19^. The scattering intensity of each sample in arbitrary scale is related to the membrane structure as

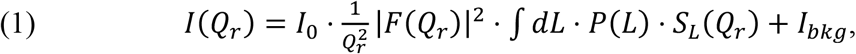

where the prefactor *1*/*Q*_*r*_ ^*2*^ is the Lorentz factor for bulk powder averaged geometry. *I*_*0*_ and *I*_*bkg*_ are the arbitrary scaling factor and the background value. *Q*_*r*_ is the scattering vector length^2,20^.

The bilayer form factor *F(Q*_*r*_) is the Fourier transformation of the bilayer SLD contrast *Δρ*_*b*_’*(z)* against the bulk value ρ_b,water_:

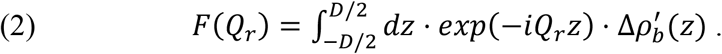

Here, a phospholipid bilayer is divided into five sub-compartments based on its SLD relative to the aqueous bulk (Figure 1B): the headgroup compartments of the two leaflets with *Δρ*_*b*_’>0, their hydrophobic tail compartments that have lower SLD than the headgroup regions, and a region between the two leaflets with the lowest SLD^9^. The maximal excess SLD of the bilayer *Δρ*_*b*_’_*max*_ = 1.

Our test systems are unextruded liposome dispersions, either with lamellar orders that show Bragg peaks in the diffractogram, or as uncorrelated bilayers without Bragg peaks. For lamellar phases, *D* is the repeat distance from the mid-point of the bulk on one side of the bilayer to the mid-point of the bulk on the other side. In the cases of uncorrelated bilayers, *D* is set to a number larger than the bilayer thickness, e.g., 100 Å. *S*_*L*_ and *P*, are structure factor of the lamellar lattice of size *L*, and the size distribution function. For uncorrelated bilayer *S*_*L*_ = 1, *P(L)* = 1 for *L* = 1 and *P* = 0 for *L* ≠1. For lamellae, the structure factor S_L_ of the bilayer lattice of size *L* is

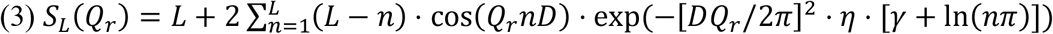

for the fluid phase of the multilamellar system^2,20^, and *γ* is the Euler-Mascheroni constant. The Caille parameter *η* ^20^, following the Pabst 2003 definition, accounts for the fluctuation of the bilayer sheet and its resulting loss of order ^2^. The normalized size distribution function *P(L)* is given by

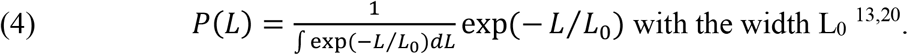

#### 2.1.2 General formula for obtaining absolute SLD profiles

The relative SLD difference *Δρ*_*b*_’*(z)* is related to the contrast of the absolute SLD profile *ρ*_*b*_*(z)* from the bulk SLD ρ_b,water_ = 9.42 × 10^−6^ Å^-2^ by

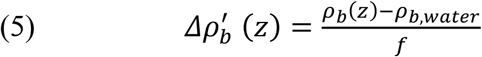

with a normalization factor

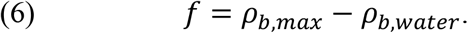

Here we will show that it is possible to calculate this normalization factor by using the physical constraint that the total scattering lengths of the hydrophobic core and hydrophilic head under the area Â of one lipid are both known, and the same normalization factor value applies over all compartments of the membrane.

Initially, integration of 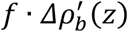 over the lipid headgroup compartment or the hydrophobic core compartment are derived from the total scattering lengths, the volume of the compartment per lipid V_head_ and V_tail_, and Â as ^21^

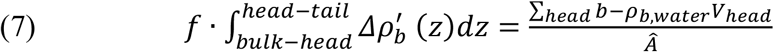

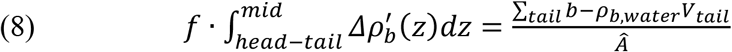

The total scattering length of the headgroup region is summed up from the electron number of the atoms constituting the headgroup, from the carbonyl-glycerol-phosphate (CGP) and the terminal groups (TER) such as choline (CHOL) or serine (SER), and hydration water (hydra):

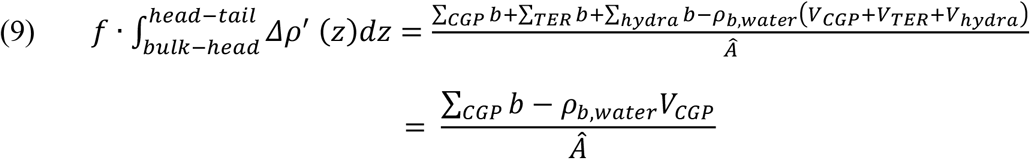

This disintegration holds regardless the SLD contrast since it is only accounting the number of electrons in the material. The TER contribution is canceled with *ρ*_*b*,*water*_ ∙ *V*_*TER*_ since its X-ray SLD is close to the bulk water, thus considered invisible ^9^.

Dividing the head and the core equation yields the solution for the area per lipid Â in the bilayer:

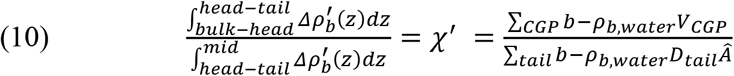

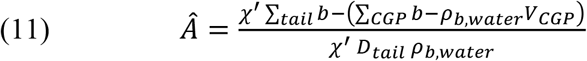

*D*_*tail*_ is the thickness of one tail leaflet, measured from the head-tail interface to the middle plane of the leaflet interface within a bilayer. The normalization factor for the absolute SLD can be obtained by using the obtained Â value as

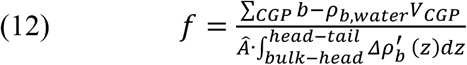

using the SLD contrast of the headgroup region. With asymmetric bilayers, the scaling factors obtained from the two leaflets should be identical, within error. The absolute SLD thus can be obtained as

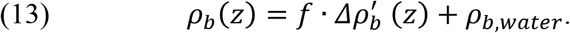

The value of *f* can equally be obtained from the contrast of tail region as

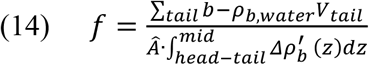

where the tail volume V_tail_ = D_tail_ ∙ Â. The integration runs through the range of D_tail_.

The calculation of *Â* and *f*, via equation 11 and 12, demands a small enough variation of the volume *V*_*CGP*_ upon change of *Â*. A variation of ±5% in *V*_*CGP*_ is acceptable, since it only attributes an uncertainty of 5% to the scaling factor *f*. Considering a typical contrast between the maximal SLD of the headgroup of a bilayer and the aqueous bulk is (12 × 10^−6^ Å^−2^) − (9.42 × 10^−6^ Å^−2^) ≈ 2.6 × 10^−6^ Å^−2 7,22^, this 5% uncertainty corresponds to only 0.05 ∙ (2.6 × 10^−6^ Å^−2^) = 0.13 × 10^−6^ Å^−2^ in the absolute SLD value. This variation has to be examined experimentally^22^, although it was often assumed to be constant for the same type of headgroup^3^.

#### 2.1.3 Determination of Reference headgroup volume from monolayers

Once the value of V_CGP_ is determined and proved to vary less than ±5%, the area Â per lipid in the bilayer and the absolute SLD can be sequentially calculated from equation 11 and 12. The approach can be practically applied under this condition, without knowing the lateral pressure in the liposome membranes. This value and its variation can be experimentally obtained from GIXOS measurements^17^ using monolayers of identical lipid compositions on the same aqueous bulk, at various lateral pressures, throughout again a conventional analysis described below^17^. In the next section we will also show that the variation V_CGP_ is indeed within 5% for a specific headgroup type that allows it to be used in equation 11 and 12.

In a GIXOS measurement, the X-ray beam is incident at the air-buffer interface at ∼85% of the critical angle of air-buffer interface, in order to enhance the monolayer scattering signal relative to the bulk scattering^23^. The GIXOS intensity is measured as a function of *Q*_*z*_, i.e. the vertical Q-component, up to 0.9 Å^-1^, at Q_xy_ → 0 Å^-1^. Beside a constant background, the measured intensity *I(Q*_*z*_) consists of three contributions as

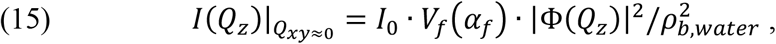

where *I*_*0*_ is an arbitrary scaling factor of the incoming flux^17,23^. *V*_*f*_, namely the Vineyard factor, is the bulk contribution (see Supplementary Information, section 1). It is a function of the critical angle of the air-buffer interface, the incident angle *α*_*i*_ and scattering angle *α*_*f* 24,25_.

The contribution from the interfacial layer *Φ(Q*_*z*_) is a Fourier transformation of the gradient *dρ*_*b*_*(z)/dz* of interfacial SLD profile *ρ*_*b*_*(z)* for a monolayer on water at an arbitrary scale. However, the SLD value of two sides, namely the superphase (air) and the subphase (water) are known to be 0 and *ρ*_*water*_. Therefore, similar to the reflectivity analysis, *dρ*_*b*_*(z)/dz* at an arbitrary scale corresponds to a unique SLD profile at absolute scale. *dρ*_*b*_*(z)/dz* and finally the absolute SLD were obtained by fitting the monolayer *I(Q*_*z*_) using a two compartment model (Figure 1B)^26^, consistent with the one leaflet of a bilayer model for SAXS: the CGP headgroup compartment with higher SLD *ρ*_*b,h,m*_ than the aqueous bulk *ρ*_*water*_, and a tail compartment with lower SLD *ρ*_*b,t,m*_ than the headgroup region. The interfaces of headgroup-bulk, tail-headgroup and air-tail are all modelled by error functions^26^. The fitting of *I(Q*_*z*_) thus yields the thickness *D*_*t,m*_, *D*_*h,m*_ of the tail and the head compartment, their SLD and the width of the error functions describing the interfaces.

Area per lipid

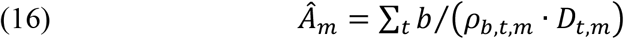

in the monolayer is then calculated from the SLD and the thickness of the tail compartment, and the total scattering length ∑_*t*_ *b* of the two hydrocarbon tails of a lipid molecule. The latter

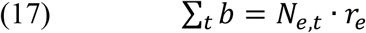

where *N*_*e,t*_ is the total electron numbers of two tails, i.e. 242e for DPPC, and 265e for POPC.

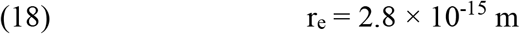

is the classical electron radius. The headgroup layer under such area contains a contribution from one CGP group. The rest of the volume has the SLD *ρ*_*b,water*_, contributed by the terminal group (choline / serine) and hydration water ^19^. Therefore

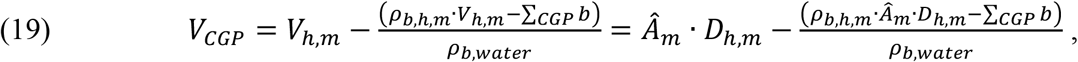

Where

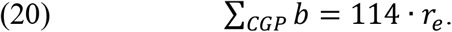

### 2.2 Experimental Section

#### Lipid stock solutions

1,2-dipalmitoyl-*sn*-glycero-3-phosphatidylcholine (DPPC), 1,2-dipalmitoyl-*sn*-glycero-3-phosphatidyl-L-serine (sodium salt, DPPS), 1-palmitoyl-2-oleoyl-PC (POPC) and POPS (sodium salt) were purchased from Avanti Polar Lipid as dry powders, and were dissolved in a mixture of chloroform / methanol (7:3 vol/vol, both from Merck KGaA) to yield four stock solutions at 1.5 mM lipid concentration. Stock solutions for binary lipid mixtures (PC:PS, 90/10 mol/mol) were obtained by mixing the PC and the PS stocks at a 90:10 volume ratio.

#### SARS-CoV-2 envelope protein purification and characterization

A SARS-CoV-2 E protein construct with N-terminal 6-His and MBP tag was derived from its SARS-CoV-1 counterpart ^1^ by introducing T55S, V56F, E69R, and G70del mutations by site-directed mutagenesis ^27^. A plasmid carrying this construct was transformed into *E. coli* BL21-CodonPlus (DE3)-RIPL (Agilent). The cells were cultivated by fed-batch method with K12 media ^2^ in a 1 L fermenter (Winpact) at 37°C. A 40% DO saturation maintained by stirring, aeration and O_2_ supplementation. Protein expression was induced with 0.5 mM IPTG at the same time as feeding was started, and cultivation continued at 18°C overnight. The culture was harvested by centrifugation at 7500×g. E protein was purified from the cell pellet by Ni-NTA chromatography and reverse-phase-HPLC as described previously ^1^. The dry E protein (M_w_ = 8541.12 g/mol) powder was then dissolved in ethanol (denatured, ≥99.8%, Carl Roth GmbH) at 1 mg/mL.

#### SAXS experiment on liposome dispersion

Liposome dispersions for SAXS measurement contains ∼23 mg/mL of total lipid in 10 mM Tris buffer with 0.05 mM EDTA at pH 7.4 with and without 0.5mol% E-protein in the lipid mixtures. Buffer solutions were first prepared and adjusted to required pH using concentrated hydrochloric acid. All buffer components were purchased from Merck KGaA. MilliQ water was provided at PETRA III (Hamburg, Germany). Lipid stock solutions of appropriate amounts were dried by rotary evaporation at 45°C, 25 mbar for 45 min. Where needed, E-protein ethanol stock solution was added to the lipid solution before drying in the rotary evaporator to a lipid:protein ratio of 200:1, equal to a 0.5mol% of E-protein in the lipid mixture. PO-lipid containing samples were rehydrated at room temperature while DP-lipid containing samples were rehydrated at 45°C, to produce a final ∼23 mg/mL lipid dispersions.

SAXS experiments were performed in mail-in mode at P12 BioSAXS beamline operated by EMBL at PETRA III (Hamburg, Germany) ^28^, using a 0.2 mm × 0.12 mm (h × v) beam at 10.0 keV, with a Pilatus 6M detector (Dectris, Switzerland) at 3 m distance. Pure water and empty capillary were measured at 22.3°C and 37.0°C, as the calibrants. Every liposome sample measurement of 10 × 0.1 s exposures was followed by at least two measurements on bare Tris buffer. All the data were first calibrated into the absolute scale [cm^-1^] using the scaling factor calculated from the water data measured at the same temperature ^29^. Background buffer data was subtracted from that of the liposome dispersions, which was then normalized by the lipid concentration. The data processing was performed with the automated pipeline DATOP with DATABSOLUTE module at the beamline ^14^.

#### GIXOS experiments on Langmuir monolayers

Grazing incidence X-ray off-specular scattering (GIXOS) experiments on Langmuir monolayers were conducted at the Langmuir GID setup at the High-Resolution Diffraction beamline P08 at PETRA III (Hamburg, Germany). Lipid monolayers were directly prepared at the Langmuir trough (30 cm × 16 cm × 0.65 cm, Riegler and Kirstein GmbH, Germany) on the sample stage, which had been filled with buffer (10 mM Tris, 0.05 mM EDTA, pH 7.4, 22.0°C). The 1.5 mM stock lipid solution was deposited dropwise, using a gastight syringe (Hamilton Company, USA), to reach 60 nmol of total lipid amount at the surface, followed by a 15 min waiting for solvent evaporation. Surface pressures of 35 mN/m, 30 mN/m and 20 mN/m were achieved by a pre-compression to 38 mN/m and an expansion to the target value, both at 40 cm^2^/min. GIXOS measurement started after the film had been relaxed at the target pressure for 15 min, while pressure was maintained and monitored. GIXOS measurement was performed with a raw mode beam of 0.20 mm × 0.07 mm (h × v) at 15 keV photon energy ^30^. The beam was deflected downwards by a quartz mirror to be incident at 0.07° at the air-buffer interface, corresponding to 85% of its critical angle. Two vertical slits of 0.25 mm and 1 mm widths were mounted 16 cm and 70 cm away from the trough centre, and offset horizontally such that only the scattered beam at 0.3° off-specular angle can pass, corresponding to Q_xy_≈0.04 Å^-1 17^. Intensity as the function of vertical scattering angle was acquired by a Pilatus 100k detector (Dectris, Switzerland) mounted after the 2^nd^ post-sample slit, with 6 × 20 s exposure time. Background scattering was suppressed by a vacuum beam path to the entrance window of the trough, a guard slit of 0.30 mm × 0.15 mm (h × v) at the entrance window, the helium atmosphere inside the trough case, and the tungsten blade of the 1^st^ post-sample slit that served as a beamstop. The vibration of the liquid surface was damped by placing a glass plate beneath the X-ray footprint area of the surface that reduced the liquid depth to less than 1 mm there ^31^. Data were reduced into I(Q_z_) with beamline provided Matlab-based tools.

#### SAXS and GIXOS data analysis

The fittings of the SAXS and GIXOS data were performed with a self-built fitting toolbox using Matlab. Their mathematics is presented in section “Theoretical background to SLD profile derivation”. The mathematical expressions of the 5-compartment model (Figure 1B) for the bilayer SLD profiles can be found in the section 3 of Supplementary Information. The description of the monolayer 2-compartment model is given in section “Theoretical background to SLD profile derivation”.

## 3 Results

The above described method was applied to liposomes of DPPC and POPC at 22°C, or two-component liposomes of DPPC with 10 mol% DPPS, and POPC with 10% POPS at 37°C, all in 10 mM Tris, 0.05 mM EDTA at pH 7.4. In addition, we also applied the method to POPC/POPS (90:10) liposomes incorporating 0.5 mol% of the envelope (E) protein of SARS-CoV-2 in the same buffer conditions at 37°C. This example was chosen because it is part of our research activities and facilitates the development of the here described method. First, we will show that the reference volume of the CGP groups determined from GIXOS experiments on monolayers of the same composition and buffer varies less than 5% at different lateral pressures. This supports the use of the volume of this particular group into the SAXS analysis for bilayers. Then we will show the absolute SLD profiles of the lipid bilayers calculated using these reference values, and their consistency with previously published data ^9,19,20^.

### 3.1 Reference headgroup volume values of typical phospholipid monolayer

Figure 2 shows the fitted GIXOS data for PC monolayers at various pressures on Tris buffer, and their SLD profiles, consistent with literature values, e.g. with a headgroup peak SLD of ∼11.8 × 10^−6^ Å^-2^ and a tail SLD of ∼8.7 × 10^−6^ Å^-2^ for condensed DPPC monolayers ^32,33^. The parameters obtained for calculating V_CGP_ are compiled in Table 1, while the full set of parameters for all systems are found in the Supplementary Information (Figure S1/S2 and Table S1). The average value of V_CGP_ for PC is almost constant at 245±10 Å^3^, and the one for 90:10 PC/PS headgroup is 252±10 Å^3^, with a variation of ±5% upon pressure change.

**Table 1.** Structure parameters of DP- and PO-monolayers from the fitting of GIXOS data. Fitting yields the thickness D_t,m_, D_h,m_ of the tail and the head compartment, their SLD ρ_b,tm_ and ρ_b,h,m_, and the width of the error function σ_a/t,m_, σ_t/h,m_ and σ_h/w,m_ for the air-tail, tail-headgroup, headgroup-water interface. V_CGP_ is the volume of the carbonyl-glycerol-phosphate group without hydration.

| monolayer | DPPC |  |  | POPC |  |  |
| --- | --- | --- | --- | --- | --- | --- |
| $\Pi$ [mN/m] | 35 | 30 | 20 | 35 | 30 | 20 |
| $\rho_{b,t,m}$ [$10^{-6} \text{ \AA}^{-2}$] | 9.26±0.07 | 9.18±0.06 | 9.16±0.06 | 8.63±0.07 | 8.53±0.07 | 8.68±0.07 |
| $D_{t,m}$ [Å] | 14.4±0.6 | 14.5±0.6 | 14.0±0.5 | 13.4±0.5 | 12.7±0.5 | 11.4±0.4 |
| $\hat{A}_m$ [Å<sup>2</sup>] | 50.8±2.2 | 51.0±2.1 | 52.8±1.9 | 64.3±2.5 | 68.7±2.8 | 75.1±2.7 |
| $V_{t,m}$ [Å<sup>3</sup>] | 732 | 738 | 741 | 861 | 870 | 855 |
| $\rho_{b,h,m}$ [$10^{-6} \text{ \AA}^{-2}$] | 11.74±0.05 | 11.68±0.05 | 11.29±0.02 | 10.98±0.04 | 10.77±0.03 | 10.50±0.03 |
| $D_{h,m}$ [Å] | 8.9±0.4 | 8.6±0.4 | 8.5±0.5 | 8.4±0.4 | 8.7±0.5 | 8.5±0.5 |
| $V_{CGP}$ [Å<sup>3</sup>] | **226±7** | **232±7** | **249±6** | **247±6** | **251±6** | **263±5** |
| monolayer | DPPC : DPPS 90:10 |  |  | POPC : POPS 90:10 |  |  |
| $\Pi$ [mN/m] | 35 | 30 | 20 | 35 | 30 | 20 |
| $\rho_{b,t,m}$ [$10^{-6} \text{ \AA}^{-2}$] | 9.00±0.06 | 9.09±0.07 | 9.04±0.07 | 8.51±0.07 | 8.60±0.07 | 8.8±0.1 |
| $D_{t,m}$ [Å] | 15.1±0.6 | 14.9±0.6 | 14.5±0.6 | 12.8±0.5 | 12.2±0.5 | 11.4±0.4 |
| $\hat{A}_m$ [Å<sup>2</sup>] | 49.8±2.0 | 50.0±2.0 | 51.7±2.2 | 68.2±2.7 | 70.6±2.9 | 74.0±2.7 |
| $V_{t,m}$ [Å<sup>3</sup>] | 753 | 746 | 750 | 872 | 863 | 841 |
| $\rho_{b,h,m}$ [$10^{-6} \text{ \AA}^{-2}$] | 11.3±0.1 | 11.24±0.04 | 10.98±0.04 | 10.83±0.04 | 10.70±0.03 | 10.50±0.03 |
| $D_{h,m}$ [Å] | 9.1±0.3 | 9.0±0.4 | 8.8±0.4 | 9.1±0.5 | 8.9±0.5 | 8.3±0.5 |
| $V_{CGP}$ [Å<sup>3</sup>] | **248±7** | **248±6** | **260±5** | **243±7** | **250±6** | **265±5** |

**Figure 2.**
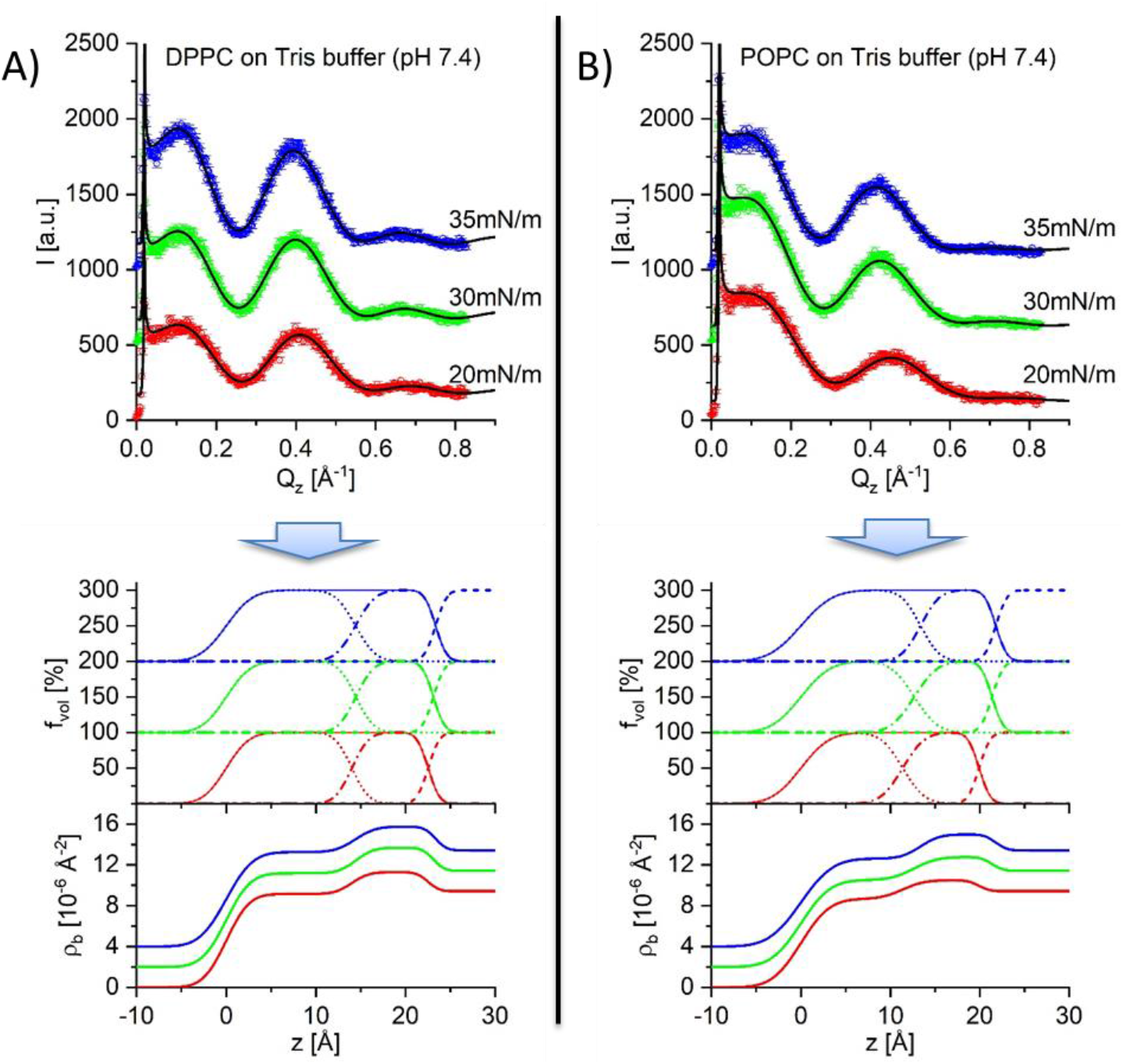
Fitting (black line) to GIXOS data (symbols) of DPPC (A) and POPC monolayers (B) at various surface pressure (offset by 500 and color-coded), and the corresponding SLD profiles to the fitting. The SLD profiles from different pressures are offset by 2, and color-coded the same as the GIXOS data. Above the SLD profile is the volume occupation f_vol_ of the tail compartment in dotted line, the headgroup compartment in dot-dashed line, the sum of the two compartments in solid line, and the bulk water in dashed line. They are offset by 100% and color-coded.

### 3.2 SLD profiles of exemplary lipid bilayers

The approach presented above was applied to finally obtain the bilayer SLD profile at absolute scale and the area per lipid in bilayers from the SAXS data of the five example systems (Figure 3). Fitting with a bilayer compartment model (Figure 1B) initially yields the relative SLD contrast profile 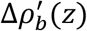 (Figure 3, inserts). The two datasets from PC-based liposome systems required the structure factor for ordered lamellae to fit the Bragg peaks and their diffuse scattering, while the three datasets from PC/PS binary liposomes were fitted with an uncorrelated bilayer model with S=1. The complete set of parameters obtained is provided in the Supplementary Information (Tables S2 and S3).

**Figure 3.**
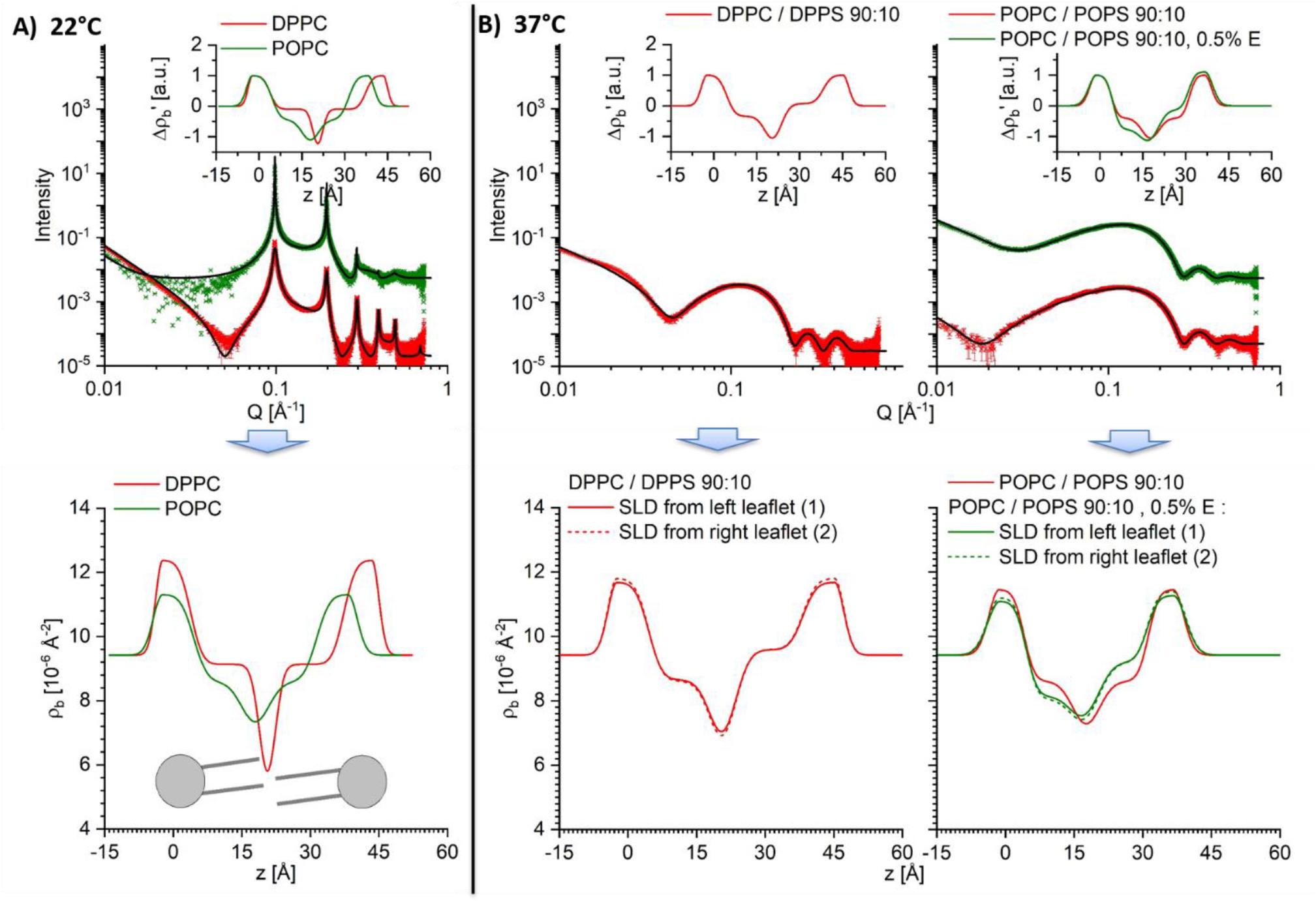
Fitted SAXS data (upper panel), SLD contrast profile (insert) and the absolute SLD profiles (lower panel) of the three PC based liposomes at 22°C, and of the two PC/PS binary liposomes at 37°C. The data from the POPC/0.5% E liposome ^#^(incorporated E protein with a 200:1 lipid-to-protein molar ratio), and from the POPC/POPS (90/10) liposomes are shown in green and both offset by 100. The SLD profiles from the two leaflets of the asymmetric bilayers are entered as different line types. The sketch of lipid molecules in the DPPC absolute SLD profile depicts the corresponding position of the headgroups, tails and the interface between the two leaflets. #: own data, not published

The 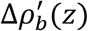 of the gel DPPC bilayer and fluid POPC bilayer are consistent with literature profiles ^6,9,15^. The shapes of the two headgroup compartments, at z=0 Å are consistent with reported values ^9,15^, with a steeper SLD drop towards the bulk and a shallower drop towards the core. The thickness D_c_ of the DPPC core region, measured as the distance between the two headgroup-tail interfaces, is 33.5 Å (Table 2), agreeing with the previously reported value of 34.2 Å ^9^. The SLD values of the two tail leaflets are constant over ∼13 Å, being close to the bulk SLD 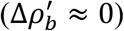. These two leaflets are separated by a narrow SLD trough 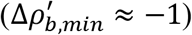, consistent with previous findings ^9^. The core thickness D_c_ of 26.9 Å in the fluid POPC bilayer, and its lower SLD than the bulk, is also consistent with published data (Table 2) ^6^. The interface between the two leaflets is wider and therefore less defined compared to the gel phase DPPC bilayer. The POPC/POPS (90/10) bilayer at 37°C has a similar structure as the POPC bilayer. However, the fitting of the data corresponding to the POPC/POPS bilayer with 0.5 mol% E protein required an asymmetric bilayer profile to account for the elevated SAXS intensity between 0.05 Å^-1^ and 0.2 Å^-1^ (Figure 3B) ^34,35^. Here, one bilayer leaflet is thicker, and has a higher SLD in both head and tail region than the other. The data of DPPC/DPPS (90/10, 37°C) also requires an asymmetric bilayer profile to provide reasonable fitting to the elevated SAXS intensity dips at ∼0.05Å^-1^ and 0.25Å^-1^. Its SLD and thickness of the two chain leaflets are slightly different (Figure 3B, Table 2).The values of headgroup area Â and the tail region volume V_tail_ per lipid were initially calculated according to Equitation 11, with the input *V*_*CGP*_ values from GIXOS measurements (242 Å^3^ for PC, and 248 Å^3^ for PC/PS), prior to the calculation of the normalization factor *f* for the absolute SLD. They were obtained for individual leaflets in the case of the asymmetric POPC/POPS/0.5% E-protein and DPPC/DPPS bilayers. This method is applicable for the POPC/POPS/0.5% E-protein bilayer since the compositional contribution of 0.5 mol% transmembrane protein to the bilayer SLD is negligible. In this regard, we included SLDs simulations to support our claim into the Supporting Information. These values together with those averaged between the two leaflets are compiled in Table 2. The *Â*-values of 45.5 Å^2^ and 64.8 Å^2^ for the reference gel DPPC and fluid POPC bilayers (Table 2) are consistent with the literature ^6,7,9^. The f-values were then obtained by Equiation 12 using the contrast of the tail leaflet (Table 2), and were finally used to obtain the absolute SLD profile (Figure 3 lower panels). In any of the asymmetric bilayers tested, the normalization factor values obtained from the two leaflets were identical (Table 2). Accordingly, a unique SLD profile was obtained independent from the choice of the leaflet (Figure 3 lower panels, solid and dashed lines).

**Table 2.** Area Â per lipid in the two leaflets (index 1 & 2) of the liposome bilayers, the SLD normalization factor f calculated from the values of Â, and the corresponding volume V_tail_ of the tail region of the lipids. The thickness D_c_ of the hydrophobic core, and the average values 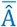 and 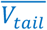 of Â and V_tail_ between the two leaflets are also entered.

| T [°C] | 22 |  | 37 |  |  |  |  |
| --- | --- | --- | --- | --- | --- | --- | --- |
| membrane | DPPC | POPC | DPPC:DPPS<br>90:10 |  | POPC:POPS<br>90:10 | POPC:POPS 90:10<br>0.5% E |  |
| V<sub>CGP</sub> [Å<sup>3</sup>] | 242 |  | 248 |  |  |  |  |
| leaflet idx | 1&2 | 1&2 | 1 | 2 | 1&2 | 1 | 2 |
| Â [Å<sup>2</sup>] | 45.5±1.2 | 64.8±1.5 | 49.0±2.5 | 43.7±1.6 | 64.3±0.8 | 74.6±3.7 | 58.3±2.4 |
| V<sub>tail</sub> [Å<sup>3</sup>] | 763±21 | 872±24 | 803±22 | 734±16 | 874±13 | 896±32 | 842.8±24 |
| **f** | **3.0±0.1** | **1.88±0.07** | **2.3±0.1** | **2.4±0.1** | **2.03±0.06** | **1.7±0.1** | **1.8±0.1** |
| D<sub>c</sub> [Å] | 33.5±0.4 | 26.9±0.4 | 33.2±0.4 |  | 27.2±0.2 | 26.5 |  |
| $\overline{\hat{A}}$ [Å<sup>2</sup>] | 45.5 | 64.8 | 46.3 | | 64.3 | 66.4 | |
| $\overline{V_{tail}}$ [Å<sup>3</sup>] | 763 | 872 | 768 | | 874 | 869.3 | |

These examples validate our stepwise approach, by the agreement of the relative SLD contrast profiles, the values of core thickness and area per lipid with the literature, and identical normalization factors obtained from the individual leaflets of one sample. Note that the fitting did not contain any constraint linking the contrast of the two leaflets.

## 4 Discussion

The intrinsic contrast variation of a lipid bilayer provides sufficient information to calculate an absolute SLD profile, once the value of the bare headgroup volume has been determined. This approach exploits the fact that the absolute SLD contrast of the headgroup and tail regions do not vary by the same factor when the area per lipid changes, due to the volume of water solvating the headgroup. The required volume of the bare headgroup can be experimentally determined from Langmuir monolayers by GIXOS measurements. The choice of Langmuir monolayers allows the use of identical membrane compositions and subphase conditions at various lateral pressures The GIXOS technique provides low background, fast measurement that can approach the monolayer form factor more than 0.9 Å^-1^ in Q_z_, such that the bare headgroup volume can be obtained accurately.

The precondition to apply the headgroup volume value, in common with Petrache’s approach, means that the dry volume of the headgroup is constant at different lateral pressures ^3^, since the lateral pressure in the bilayer is unknown. The experimentally obtained value here varies by 5% between 20 mN/m and 35 mN/m, and between the fluid and the gel phase (Table 1). However, the impact of this variation is negligible since it only generates an uncertainty at 5% of the difference between the headgroup maximal SLD and the bulk water:

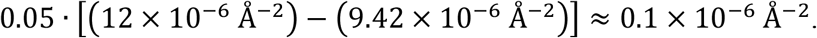

Our five examples demonstrate the application of this method to both symmetric and asymmetric bilayers. The method was first validated by the literature-consistent values of area per lipid for the reference DPPC and POPC liposomes (Table 2) ^6,7,9^. The theoretical derivation was further supported by the identical SLD profile obtained from the two different leaflets of the asymmetric bilayers (Figure 3), without providing constraints that may couple the SLD of the two leaflets in SAXS fitting. We indeed notice that for the DPPC system, SLD scaling previously obtained is larger, which leads to a greater maximal SLD of the headgroup, up to 13.8 × 10^−6^ Å^-2^, and a lower minimal SLD value down to 0.45 × 10^−6^ Å^-2 6,9,15^. This apparent discrepancy is due to the different CGP headgroup volumes applied, i.e a recent value of 178 Å^3^ for gel phase DPPC ^9^, 213 Å^3^ for fluid DMPC and POPC ^6,7,15^, both obtained from simulations. Applying these two values to our calculation yields identical SLD profiles as theirs (Supplementary Information, Figure S3), which further serves to validate our approach. Nevertheless, the volume of the bare headgroup may indeed vary upon the change of physico-chemical conditions ^22^. We therefore recommend using experimentally determined volumes of the CGP group at the closest conditions possible, such as in the method here reported. We are confident in using our values of 245 Å^3^ for PC, and 252 Å^3^ for PC/PS (90:10), which were determined with the same lipid compositions in the same buffer, using monolayer SLD profiles consistent with data published from similar techniques ^32,33,36,37^.

Moreover, the SLD profiles obtained provide an opportunity to investigate the relationship between a fully hydrated bilayer and an air-buffer interface monolayer. Figure 4 shows a comparison between the liposome bilayer cross sections and those of their equivalent monolayers at 35 mN/m (22°C), which is considered to correspond to lateral pressure in a bilayer ^38,39^. Although it is not possible to compile the SLD of a bilayer as the result of an imposition of two SLD of a monolayer, it is interesting to note that the two fluid PO-bilayers resemble the superimposed mirrored SLD profiles from two monolayers of the same lipid composition (Figure 4, left), when their headgroup centroids are superimposed over those of the two bilayer headgroups. At this position, the CH_3_-terminal interfaces of the two monolayer tail compartments are slightly separated at a distance of 0.9 Å (Figure 4, vertical dashed lines). This separation mimics precisely the “low SLD trough” as observed at the centre of the PO-liposome bilayers. A similar phenomenon was previously observed for myelin lipid membranes ^40^. For the gel DP-case, the superimposed SLD profiles qualitatively resemble the profile of the DP-liposome bilayers, although the details differ. Note that the superposition of two monolayers does not consider the bilayer asymmetry and we do not recommend to use SLD profiles of monolayers to infer the SLD of a bilayer. Still, the headgroup compartments have a similar shape, width and SLD. The average SLD of the two tail compartments of the liposome bilayer equals the SLD of the monolayer tail region. However, superimposing the monolayer and bilayer headgroup centroids yields a larger separation of 3.8 Å between the CH_3_-terminal interfaces of the DP-monolayers, compared to the PO-case. This larger separation is qualitatively consistent with the deeper “SLD trough” between the DP-leaflets. Thus, the experimentally observed trough is not as deep as the reconstructed one from mirror image monolayers, suggesting that the interface between the two bilayer leaflets bilayer is less defined than the monolayer tail-air interface. Beside the direct observation we reported from our analysis, this discrepancy at interfaces between monolayer and bilayer SLDs is a further confirmation that structural data from monolayer studies need to be carefully considered when applied to bilayer analysis. Also, for this reason, the presented method aims to justify theoretically and experimentally the reasoning behind the use of GIXOS as aid to SAXS analyses.

**Figure 4.**
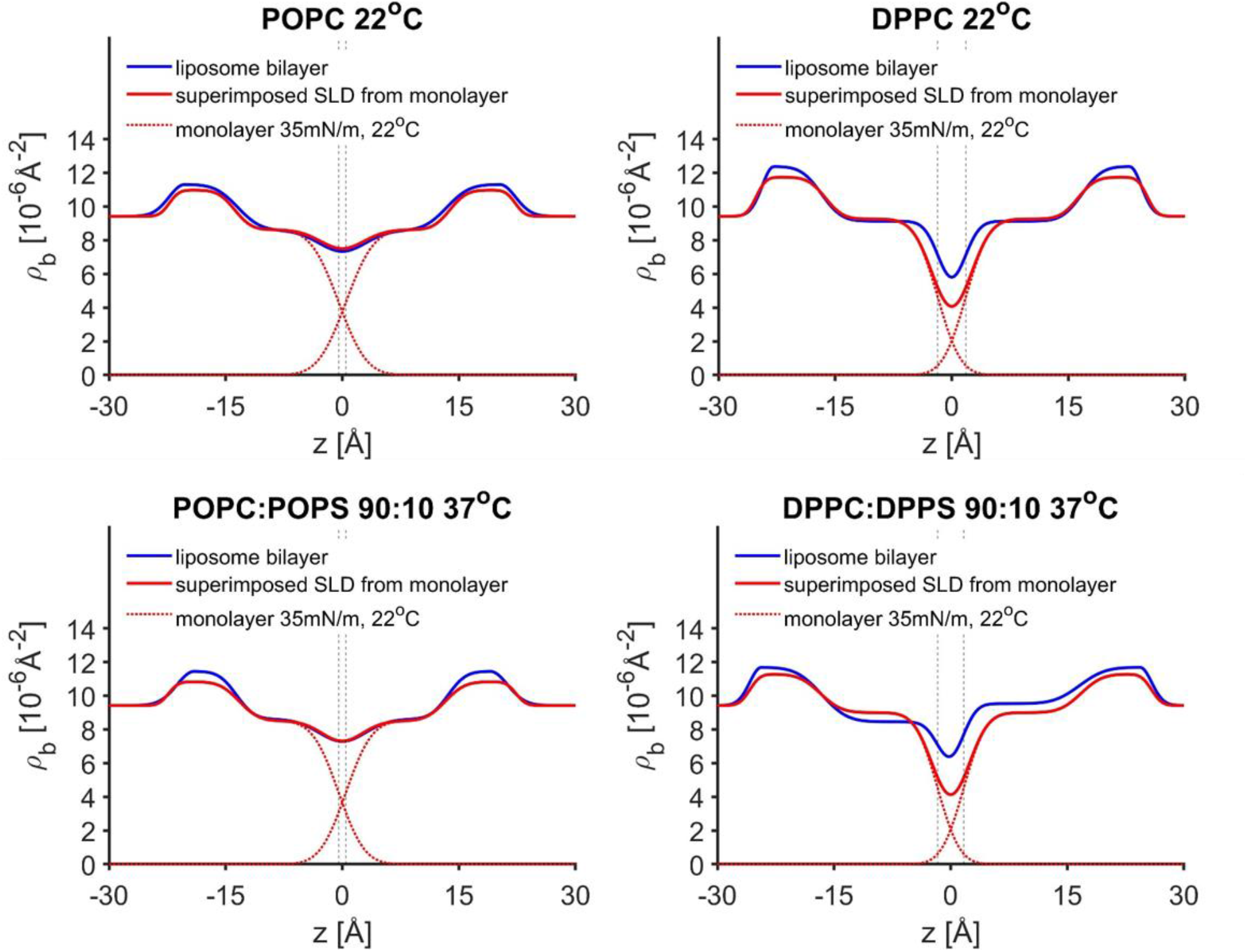
Absolute SLD profiles of the bilayers in liposomes versus the superimposed SLD profile from two mirrored monolayers of the same kind, at 35mN/m, 22°C. The monolayers are placed such that their headgroup centroids coincide with the centroids of the two bilayer headgroup compartments. The monolayer SLD profiles are also entered, as dashed red lines. The resulting position of the CH_3_ terminal interface of the monolayer tail region is depicted by dashed vertical line.

## 5 Conclusion

In summary, we present a purely experimental approach with X-ray measurements to obtain the area per lipid and thus the absolute SLD profile of liposome membranes. Beside the SAXS data, the only input required is the headgroup volume, which we suggest to measure at identical lipid composition and subphase conditions. e.g., from GIXOS experiments. The method is validated by selected reference phospholipid liposome systems. In this article, we also provide the full set of equations needed and the criteria of the data for such calculations. This approach is easily accessible as GIXOS experiments on Langmuir monolayers are routinely performed at major synchrotron facilities ^17,41^, whereas suitably high-quality solution SAXS experiments are achievable with both synchrotron and lab sources.

(SLD): Scattering length density
(SAXS): small X-ray scattering
(GIXOS): grazing incidence off-specular scattering
(E): envelope protein
(SARS-CoV-2): the severe acute respiratory syndrome coronavirus 2
(CGP): carbonyl-glycerolphosphate
(TER): terminal groups
(CHOL): choline
(SER): serine

## 6 Acknowledgements

We acknowledge DESY (Hamburg, Germany), a member of the Helmholtz Association HGF, for the provision of experimental facilities. Parts of this research were carried out at PETRA III and we would like to thank Dr. F. Bertram and Mr. R. Kirchhof for assistance in using beamline P08, Dr. M. Lippmann, Mrs. A. Ciobanu for assistance in using the chemistry laboratory. Beamtime was allocated for fast track corona proposal P-20010207 and proposal H-20010161. The synchrotron SAXS data was collected at beamline P12 operated by EMBL Hamburg at PETRA III for the proposal SAXS-1095. The research leading to this result has been supported by the project CALIPSOplus under the Grant Agreement 730872 from the EU Framework Programme for Research and Innovation HORIZON 2020. CS thanks Prof. B. Klösgen (SDU, Odense, Denmark), Prof. J. F. Nagle and Prof. S. Tristram-Nagle (CMU, Pittsburgh, USA) for the fruitful discussion on the structural analysis of lipid membranes. JT and WS thank the Singapore Ministry of Education (MOE) Tier 1 thematic grant RT13/19.

## 7 Author contributions

CS, GB, RH, CW developed the methodology, performed the GIXOS experiments, prepared the samples for the SAXS experiments, analysed all the data, did the major manuscript writing and conceptualized the research. CS also implemented the fitting procedure. AK did the mail-in SAXS measurement at P12. JT and WS provided the SARS-CoV-2 E protein and the protocol to prepare the liposomes with the protein and also contributed to the writing and conceptualization.

