## Supplemental Information for "Absolute scattering length density profile of liposome bilayers obtained by SAXS combined with GIXOS - a tool to determine model biomembrane structure"

Supplementary information

### Absolute scattering length density profile of lipid bilayers of liposome obtained by solution-SAXS combined with GIXOS on Langmuir monolayers

Chen Shen, Christian Woelk, Alexey G. Kikhney, Jaume Torres, Wahyu Surya, Richard D. Harvey, Gianluca Bello

#### 1 Vineyard peak contribution to GIXOS intensity

The background free GIXOS intensity $I(Q_z)$ is proportional to the product of Vineyard factor $V_f$ that is related to the scattering length density (SLD) difference between air and water, and the structure factor $\Phi(Q_z)$ of the interfacial layer [1,2]

$$I(Q_z)|_{Q_{xy}\approx 0} \propto I_0 \cdot V_f(\alpha_f) \cdot |\Phi(Q_z)|^2/\rho_{b,water}^2 \quad . \qquad \text{eq. S1}$$

$V_f$ scales the scattered intensity as a function of the outgoing angle $\alpha_f$ , roughly from zero to a factor of two when $\alpha_f$ increases to the critical angle of the interface, and then rapidly drop to 1 [3]. It can be expressed mathematically as [4,5]:

$$V_f(\alpha_f) = T(\alpha_f) \cdot \frac{\Lambda(\alpha_f)}{\Lambda_0} \quad , \qquad \text{eq. S2}$$

where T is the Fresnel transmittivity from the air-water interface with absorption contribution:

$$T(\alpha_f) = \left| \frac{2\left(\alpha_f/\alpha_c\right)}{\alpha_f/\alpha_c + \sqrt{\left(\alpha_f/\alpha_c\right)^2 - 1 - \frac{2\beta}{\alpha_c^2} i}} \right|^2 \quad .$$

$\beta = 10^{-9}$ and $\alpha_c = 0.081°$ are the imaginary part of the refractive index and the critical angle, respectively, both of water for 15 keV X-ray beam. The calculation of the transmittivity also considers the variation of $\alpha_f$ due to the ~70 cm footprint.

Λ is the penetration path of the incident and the scattered beam:

$$\Lambda(\alpha_f) = \frac{1}{k} \cdot \frac{1}{l_i + l_f}$$

$$l_{i,f} = \frac{1}{\sqrt{2}} \cdot \sqrt{\left(\sin^2 \alpha_c - \sin^2 \alpha_{i,f}\right) + \left[\left(\sin^2 \alpha_{i,f} - \sin^2 \alpha_c\right)^2 + (2\beta)^2\right]^{\frac{1}{2}}}$$

where k is the wavenumber of X-ray beam. The constant $\Lambda_0 = (k \cdot \ell_i)^{-1}$ is Λ at large $\alpha_f$ limit.

#### 2 GIXOS results and the structural parameters of the monolayers

The monolayer GIXOS data were analyzed using a 2-slab compartment model as presented in the main text (Figure 1b) [6]. Figure S1 shows the fitted GIXOS data from all the monolayers measured. Figure S2 shows the corresponding SLD profiles obtained from the analysis using the 2-slab compartment model (main text, Figure 1b), and the volume occupation of each slab and the subphase. The complete set of the obtained structural parameters that describes the SLD profile is compiled into Table S1.

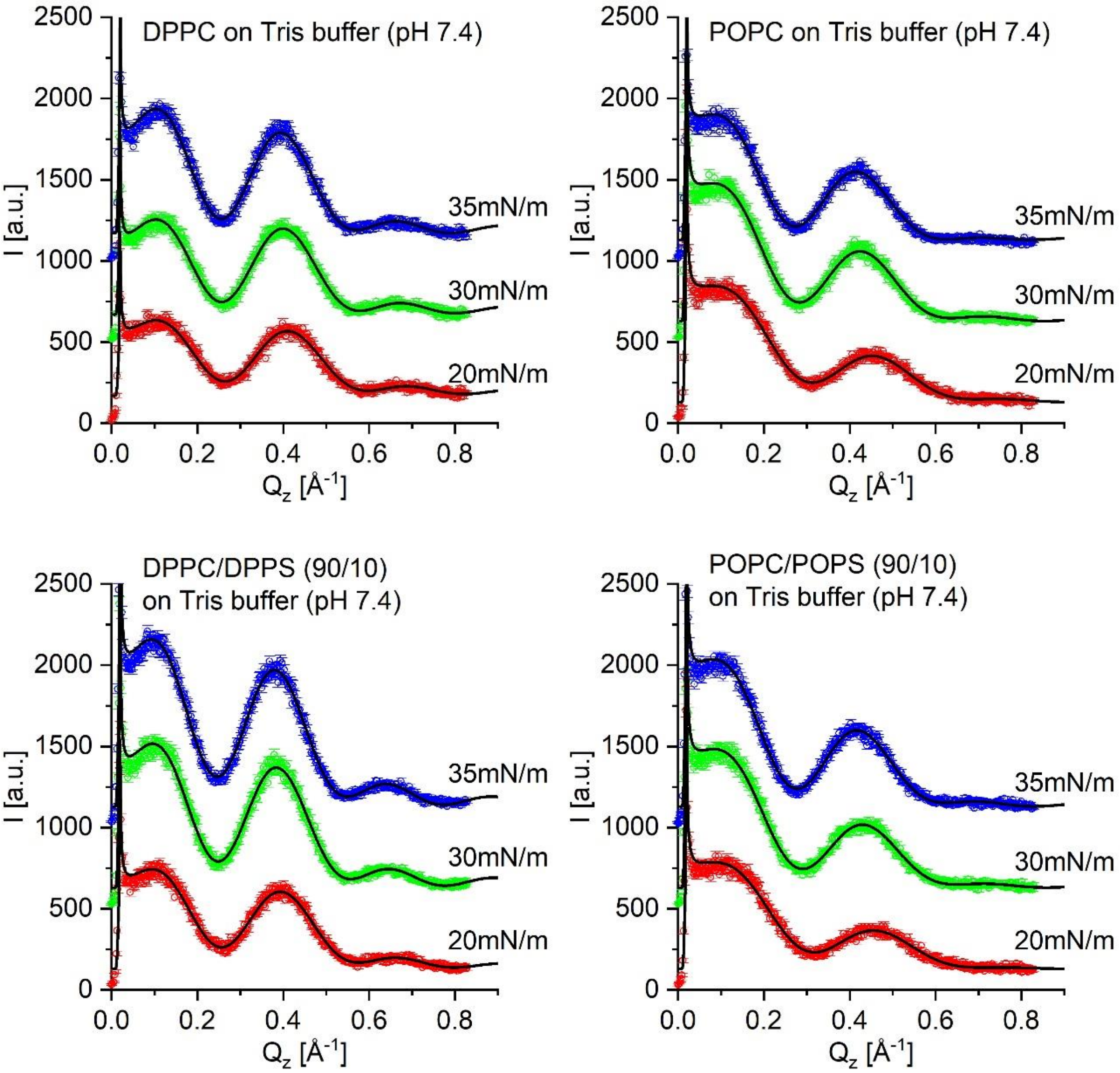


*Figure S1 GIXOS data from 4 monolayers of different composition, all on 10mM Tris buffer at pH 7.4, at 22°C, together with the fitting (black line) using 2-slab compartment model. Data from different pressure values are offset by 500 and color-coded.*

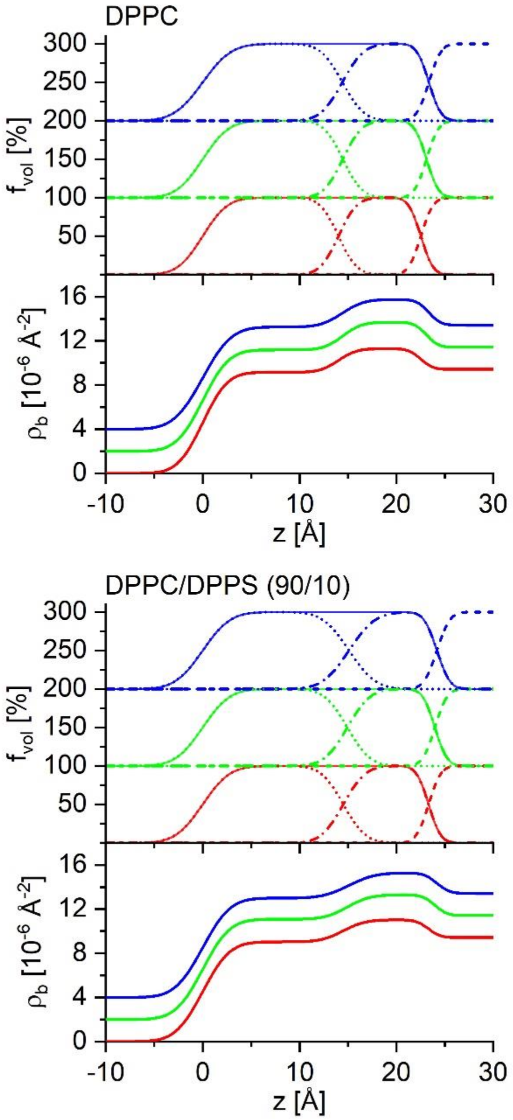


*Figure S2 SLD profiles and the volume occupation profiles of the tail, the headgroup and the subphase compartment, that correspond to the fittings of the GIXOS data in Figure S1. Profiles for the monolayer at 20mN/m, 30mN/m and 35mN/m are color-coded as before, and offset (by 2 in SLD, and by 100% in the volume occupation profiles).)*

*Table S1 Complete set of the structural parameters of the DP- and PO-monolayers obtained from the fitting of GIXOS data using 2-slab compartment model. Subscripts follow the definition in the main text: w for water subphase, h for headgroup compartment, t for tail compartment, a for air, and m for monolayer. $\rho_{b,i,m}$, $D_{i,m}$ and $\sigma_{i/j,m}$ are the SLD and the thickness of the compartment-i of the monolayer, and the width (standard deviation) of the error function that models the interface between the compartment-i and compartment-j, respectively.*
**: uncertainty of $\sigma_{h/w,m}$ are mostly below 0.01Å, except for DPPC:DPPS (90:10) binary monolayer at 35mN/m*

| monolayer | DPPC | | | POPC | | |
|---|---|---|---|---|---|---|
| Π [mN/m] | 35 | 30 | 20 | 35 | 30 | 20 |
| $\sigma_{a/t,m}$ [Å] | 2.3±0.1 | 2.1±0.1 | 2.03±0.05 | 2.8±0.1 | 2.6±0.1 | 2.5±0.1 |
| $\rho_{b,t,m}$ [$10^{-6}$ Å$^{-2}$] | 9.26±0.07 | 9.18±0.06 | 9.16±0.06 | 8.63±0.07 | 8.53±0.07 | 8.68±0.07 |
| $D_{t,m}$ [Å] | 14.4±0.6 | 14.5±0.6 | 14.0±0.5 | 13.4±0.5 | 12.7±0.5 | 11.4±0.4 |
| $\sigma_{t/h,m}$ [Å] | 1.77±0.05 | 1.57±0.05 | 1.50±0.04 | 1.70±0.06 | 2.06±0.07 | 1.9±0.1 |
| $\rho_{b,h,m}$ [$10^{-6}$ Å$^{-2}$] | 11.74±0.05 | 11.68±0.05 | 11.29±0.02 | 10.98±0.04 | 10.77±0.03 | 10.50±0.03 |
| $D_{h,m}$ [Å] | 8.9±0.4 | 8.6±0.4 | 8.5±0.5 | 8.4±0.4 | 8.7±0.5 | 8.5±0.5 |
| $\sigma_{h/w,m}$ [Å] | 1.0 * | 1.0 | 1.0 | 1.0 | 1.0 | 1.0 |
| monolayer | DPPC : DPPS 90:10 | | | POPC : POPS 90:10 | | |
| Π [mN/m] | 35 | 30 | 20 | 35 | 30 | 20 |
| $\sigma_{a/t,m}$ [Å] | 2.2±0.1 | 2.2±0.1 | 2.2±0.1 | 2.6±0.1 | 2.6±0.1 | 2.7±0.1 |
| $\rho_{b,t,m}$ [$10^{-6}$ Å$^{-2}$] | 9.00±0.06 | 9.09±0.07 | 9.04±0.07 | 8.51±0.07 | 8.60±0.07 | 8.8±0.1 |
| $D_{t,m}$ [Å] | 15.1±0.6 | 14.9±0.6 | 14.5±0.6 | 12.8±0.5 | 12.2±0.5 | 11.4±0.4 |
| $\sigma_{t/h,m}$ [Å] | 2.1±0.1 | 1.88±0.06 | 1.81±0.07 | 2.0±0.1 | 2.0±0.1 | 1.6±0.1 |
| $\rho_{b,h,m}$ [$10^{-6}$ Å$^{-2}$] | 11.3±0.1 | 11.24±0.04 | 10.98±0.04 | 10.83±0.04 | 10.70±0.03 | 10.50±0.03 |
| $D_{h,m}$ [Å] | 9.1±0.3 | 9.0±0.4 | 8.8±0.4 | 9.1±0.5 | 8.9±0.5 | 8.3±0.5 |
| $\sigma_{h/w,m}$ [Å] | 1.1±0.2 | 1.0 | 1.0 | 1.0 | 1.0 | 1.0 |

### 3 Mathematical model and structural parameters of the bilayers obtained from SAXS fitting

SAXS data from liposome samples were fitted by a five-compartment model (Figure 1b, main text), that consists of headgroup slabs (n=1, 5) toward the aqueous bulk at both sides of the bilayer, and two tail slabs (n=2, 4) in between the headgroup slabs that are separated by a low SLD leaflet interface slab (n=3). The headgroup slab is mathematically described by the combination of an error function for its interface towards the tail compartment, and a half-Gaussian peak towards the bulk side, such as:

$$\Delta\rho_b'(z) = \begin{cases} \Delta\rho_{b,h}' \cdot \left[0.5 + 0.5 \cdot \mathrm{erf}\left(\frac{z-(z_0-3\sigma_{h/t})}{\sqrt{2}\cdot\sigma_{h/t}}\right)\right] & , z < z_0 \\ \Delta\rho_{b,h}' \cdot \exp\left(\frac{-(z-z_0)^2}{2\sigma_{w/h}^2}\right) & , z \geq z_0 \end{cases}$$

for the headgroup where the bulk is on the larger z side. Here, $z_0$ is the position with the maximal SLD contrast $\Delta\rho_{b,h}'$. $\sigma_{h/t}$ and $\sigma_{w/h}$ are the standard deviation of the error function modelling the transition towards the tail region, and the standard deviation of the Gaussian function modelling the transition towards the bulk, respectively. The centre of the error function therefore is $3\sigma_{h/t}$ away from $z_0$, such that the SLD reaches to $\Delta\rho_{b,h}'$ at $z_0$ when approaching from the tail side.

The three core compartments of the symmetric bilayer are described all together, in a simplified fashion, by one slab with a fundamental contrast value $\Delta\rho_{b,t}'$, and error functions on the sides towards the two headgroups. The leaflet interface at the centre is modelled by a further subtraction of a Gaussian function with amplitude $\Delta\rho_{b,mid}'<0$ at the middle of this slab:

$$\Delta\rho_b'(z) = \Delta\rho_{b,t}' \cdot \left[0.5 \cdot \mathrm{erf}\left(\frac{z-z_0}{\sqrt{2}\cdot\sigma_{h/t,1}}\right) - 0.5 \cdot \mathrm{erf}\left(\frac{z-(z_0+D_c)}{\sqrt{2}\cdot\sigma_{h/t,2}}\right)\right] + \Delta\rho_{b,mid}' \cdot \exp\left(\frac{-(z-(z_0+D_c/2))^2}{2\sigma_{mid}^2}\right) .$$

Here $z_0$ and $D_c$ are the position of the first headgroup/tail interface (small z side), and the thickness of the slab. The position of the $2^{nd}$ interface (tail/head) thus is at $z_0$+$D_c$. $\sigma_{h/t,1}$=$\sigma_{h/t,2}$ are the standard deviation of the two error functions, and must be equal to the corresponding value applied to the headgroup function. $\sigma_{mid}$ and $z_0 + D_c/2$ are the Gaussian standard deviation and the center of the Gaussian function for the leaflet interface, respectively.

The values obtained for the two symmetric bilayer samples are compiled into Table S2.

*Table S2 Structural parameters for the symmetric bilayers, as obtained from the SAXS fitting using 5-compartment model. The rows belong to the head, tail and the leaflet interface are color-coded as blue, red and orange, respectively. The Caille parameter η, lamellar repeat distance D, lamellar lattice size $L_0$ for the structure factor are also entered, together with the intensity fraction $f_{diff}$ from the lamellae diffraction. $I_0$ is an arbitrary scaling factor for the intensity. † represents the fixed input value during the fitting.*

| bilayer | DPPC | POPC | POPC:POPS 90/10 |
|---|---|---|---|
| T [°C] | 22 | | 37 |
| $\sigma_{w/h}$ [Å] | 1.5±0.2 | 2.00±0.05 | 2.24±0.03 |
| $\Delta\rho'_{b,h}$ [a.u.] | 1 † | 1 † | 1 † |
| $\sigma_{h/t}$ [Å] | 2.04±0.06 | 2.28±0.04 | 1.80±0.02 |
| $D_c$ [Å] | 33.5±0.4 | 26.9±0.4 | 27.2±0.2 |
| $\Delta\rho'_{b,t}$ [a.u.] | -0.10±0.01 | -0.44±0.01 | -0.400±0.003 |
| $\Delta\rho'_{b,mid}$ [a.u.] | -1.13±0.05 | -0.66±0.03 | -0.65±0.01 |
| $\sigma_{mid}$ [Å] | 1.8±0.1 | 3.0±0.1 | 3.00±0.05 |
| η | 0.009±0.005 | 0.049±0.006 | -- |
| D [Å] | 63.4 | 63.8 | -- |
| $L_0$ | 9±2 | 95±1 | -- |
| $f_{diff}$ [%] | 57±16 | 100 | 0 † |
| $I_0$ [a.u.] | 0.39±0.03 | 0.014±0.001 | 1.0±0.2 |

Asymmetric bilayer models use the same headgroup shape, and a slightly modified mathematical description on the core region. The three compartments are described by three slabs, all with two error function sides. Two of them represent the tail leaflets, with different SLD $\Delta\rho'_{b,2}$, $\Delta\rho'_{b,4}$ and thickness $D_{t,2}$, $D_{t,4}$. A rather thin slab with a fixed $\Delta\rho'_{b,3}$= -2 is inserted in between the two tail slabs, describing the leaflet interface region with low SLD. With this fixed contrast, the combination of slab thickness $D_3$ and the error function width $\sigma_{3/2} = \sigma_{4/3}$ was sufficient to describe any shape of this low SLD region with varying minimum. In addition, the SLD maxima of the two headgroups $\Delta\rho'_{b,1}$ and $\Delta\rho'_{b,5}$ may allow to differ when needed.

The obtained parameter values are compiled into Table S3.

*Table S3 Structural parameters for the asymmetric bilayers, as obtained from the SAXS fitting using 5-compartment model. The rows belong to the head, tail and the leaflet interface are color-coded as blue, red and orange, respectively. The parameters for the lamellae diffraction are defined as in Table S2. † represents the fixed input value during the fitting. * represents the headgroup slab 5 parameters that are set to be symmetric with the headgroup slab 1.*

| bilayer | DPPC : DPPS 90/10 | POPC : POPS 90/10 0.5% E |
|---|---|---|
| T [$^o$C] | 37 | 37 |
| $\sigma_{w/5}$ [Å] | 1.5 * | 2.7 * |
| $\Delta\rho'_{b,5}$ [a.u.] | 1 * | 1.11±0.03 |
| $\sigma_{5/4}$ [Å] | 2.6 * | 1.9 * |
| $D_4$ [Å] | 16.6±0.2 | 11.6±0.3 |
| $\Delta\rho'_{b,4}$ [a.u.] | 0.05±0.02 | -0.12±0.03 |
| $D_3$ [Å] | 2.7±0.2 | 3.3±0.2 |
| $\Delta\rho'_{b,3}$ [a.u.] | -2 † | -2 † |
| $\sigma_{4/3}$, $\sigma_{3/2}$ [Å] | 1.5±1.3 | 3±2 |
| $D_2$ [Å] | 16.6±0.2 | 11.6±0.3 |
| $\Delta\rho'_{b,2}$ [a.u.] | -0.43±0.02 | -0.76±0.03 |
| $\sigma_{2/1}$ [Å] | 2.6±0.1 | 1.90±0.05 |
| $\Delta\rho'_{b,1}$ [a.u.] | 1 † | 1 † |
| $\sigma_{1/w}$ [Å] | 1.5±0.2 | 2.7±0.1 |
| $I_0$ [a.u.] | 0.9±0.1 | 0.7±0.1 |

#### 4 Influence of the value of $V_{CGP}$

Figure S3 shows the impact of the headgroup volume $V_{CGP}$ on the obtained absolute SLD profiles. We applied $V_{CGP}$ = 178 Å$^3$ used in Nagle's DPPC result [7] into our calculation for the gel phase DPPC (22$^o$C), and $V_{CGP}$ = 213 Å$^3$ used by Klauda [8] and Heftberger [6] to calculate the profile of the fluid POPC (22$^o$C). The SLD profiles yielded are converted into the electron density profile (EDP) and entered as red lines, to be compared with the EDP converted from the SLD profiles that we obtained (black lines). The plot scale is set to the same as in previous publications [7-9] for the ease of comparison. The smaller $V_{CGP}$ input values result in a larger electron density contrast amplitude from the bulk value (0.33e/Å$^3$), and reproduce the EDP as in previous publications. The influence is more pronounced in the EDP of DPPC, while the result on POPC from us is consistent with the literature EDP. This proves the validity of our approach of using the internal SLD contrast, and also the importance to determine the precise value of the CGP headgroup volume.

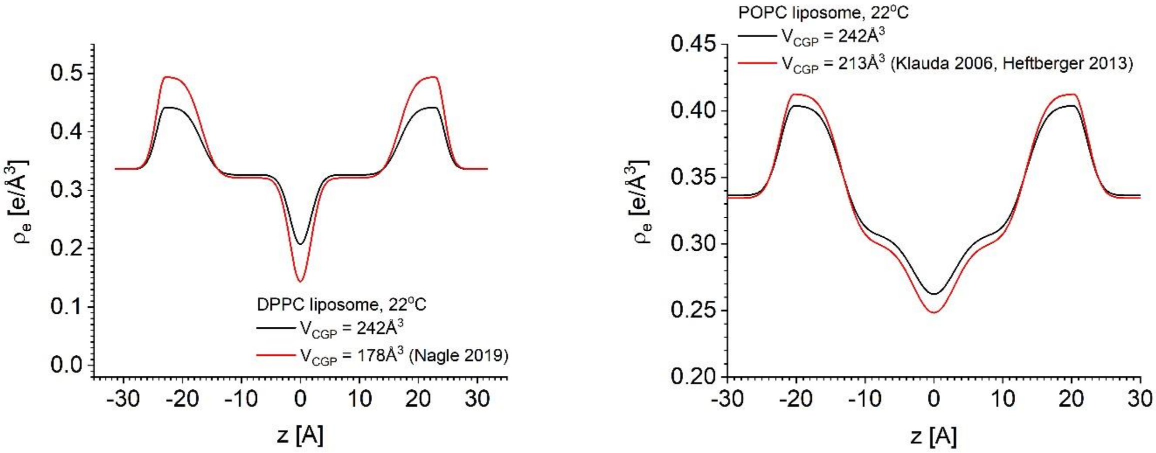


*Figure S3 Electron density profiles obtained by using $V_{CGP}$ = 242Å$^3$ from the GIXOS measurements, and by using the literature values: 178Å$^3$ from for gel phase DPPC [7], and 213Å$^3$ for fluid membranes (POPC, fluid DMPC) [8,9].The scale is adjusted to be the same as in the corresponding literatures for ease of comparison. Note the conversion between the electron density and scattering length density by a factor of classical electron radius.*

#### 5 Estimation of SLD and area contribution of the peptide to the bilayer

This simulation shows the neglectable contribution of the 0.5 mol% peptide inserted into the bilayer.

**Area occupied by the helix of the transmembrane domain in a lipid bilayer:**
approximately 60 Å$^2$ for a lipid and alpha helix d~10 Å (see figure below), $\pi r^2 \approx 80$ Å$^2$,

with a protein content of 0.5 mol % in a bilayer translates into 1.3 area% occupied by the helix.

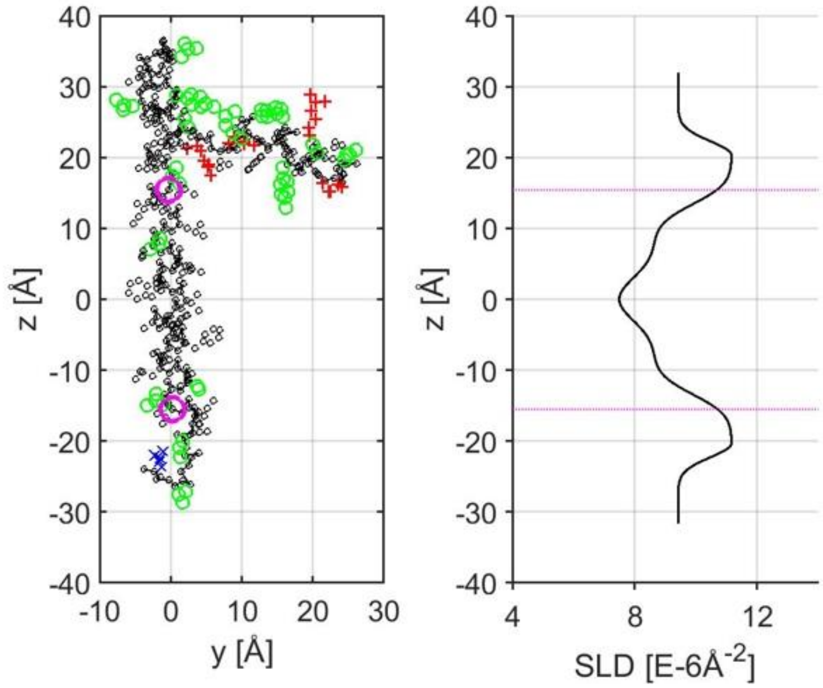


*Figure S4. Side view of the E protein structure (left) and SLD profile of the bilayer of a POPC liposome (22°C) (right). The transmembrane region of E protein (residues 17 to 37, purple circles) is aligned to the hydrophobic core of the bilayer. The position of the α-carbon in these residues is indicated relative to the SLD profile (purple lines). Black circles represent the amino acid backbone and apolar residues. The red plus sign, blue cross and green circles represent polar residues with positive and negative charges, and neutral polar residues, respectively.*

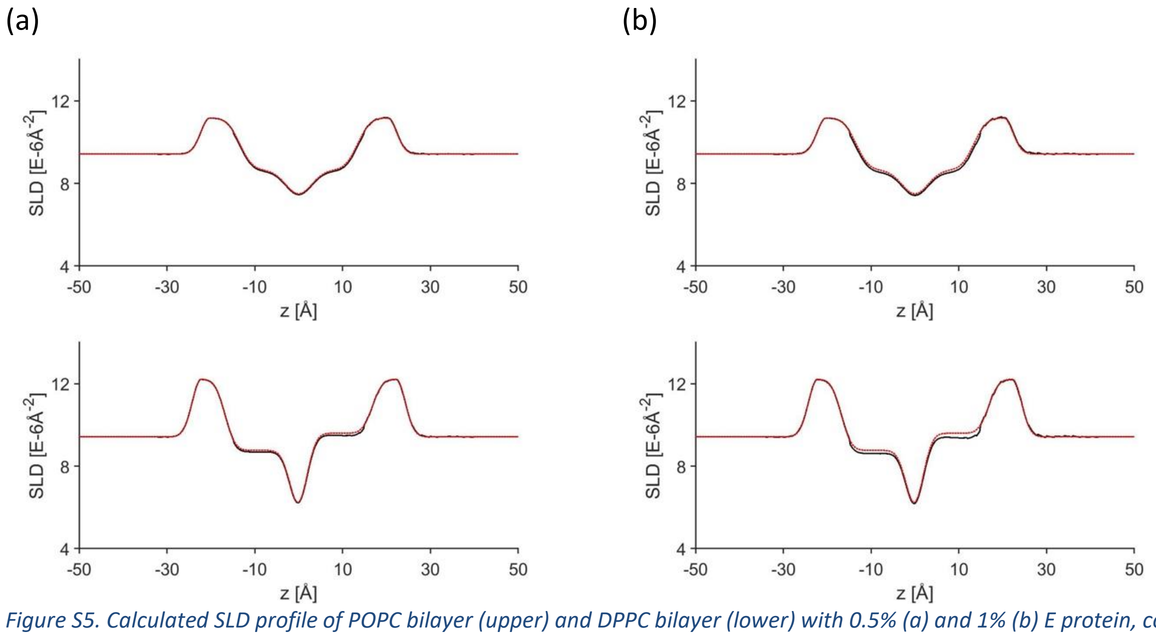


*Figure S5. Calculated SLD profile of POPC bilayer (upper) and DPPC bilayer (lower) with 0.5% (a) and 1% (b) E protein, compared to SLD of pure bilayers (red dashed).*

**Number of peptides in one 100 nm diameter vesicle:**

$$N_{pep} = \frac{4\pi\left(500\text{Å}\right)^2}{60\text{Å}^2} \cdot 0.5\% \approx 260$$

Membrane area around each peptide on average

$$\tilde{A} = \frac{1}{0.5\%} \cdot 65\ \text{Å}^2 + 80\ \text{Å}^2 \approx 13000\ \text{Å}^2 = 130\ \text{nm}^2$$
